# From distyly to stigma-height dimorphism: phylogenetic insights into the evolutionary breakdown of distyly in *Jasminum*

**DOI:** 10.64898/2026.08.16.744856

**Authors:** Shiqiao Chen, Kai Zhang, Jianyu Zhang, Yu Zhang, Xia Peng, Miaomiao Shi, Xiangping Wang, Shijin Li, Zhonghui Ma, Tieyao Tu, Zhongtao Zhao, Dianxiang Zhang

**Affiliations:** Laboratory of Plant Resources Conservation and Sustainable Utilization, South China Botanical Garden, Chinese Academy of Sciences, Guangzhou 510650, China; Ministry of Education Key Laboratory for Ecology of Tropical Islands, Key Laboratory of Tropical Animal and Plant Ecology of Hainan Province, College of Life Sciences, Hainan Normal University, Haikou 571158, China; College of Agriculture, National Demonstration Center for Experimental Plant Science Education, Guangxi University, Nanning 530004, Guangxi, China; Applied Research Center for Tropical Plant Conservation & Southeast Asia Biodiversity Research Institute, Xishuangbanna Tropical Botanical Garden, Chinese Academy of Sciences, Mengla 666303, China

**Author notes:** **Correspondence:** Zhongtao Zhao.

**Keywords:** ancillary polymorphisms, distyly, heteromorphic incompatibility, *Jasminum*, stigma-height dimorphism

## Abstract

**Background and Aims:** Distylous polymorphisms (distyly) are adaptations for plants to improve efficiency of cross-pollination and reduce pollen wastage. Theoretical models suggest that distyly evolves from stylar monomorphism via an intermediate stage of stigma-height dimorphism (SHD), however, this evolutionary scenario is only observed in few distylous lineages. The genus *Jasminum* have many species exhibit either distyly or SHD, providing an ideal opportunity to test models of the evolution of distyly. Focusing on the evolution of distyly, we investigated floral morphs and evaluated the occurrence of distyly and SHD in *Jasminum* using phylogenetic reconstruction and morphological analysis.

**Methods:** We investigated floral morphs and evaluated the occurrence of distyly and stigma-height dimorphism through morphology observation. To perform phylogenetic analysis, we sequenced plastomes of forty species from *Jasminum* and *Chrysojasminum* using Illumina next generation sequencing, and then constructed phylogenetic tree using maximum likelihood method and Bayesian inference. Based on the phylogenetic tree, we inferred the ancestral floral type through ancestral reconstruction.

**Key Results:** Our results suggest that the distyly originated in the common ancestor of *Jasminum* and *Chrysojasminum*. SHD occurs in several *Jasminum* species scattered in different clades of the phylogenetic tree, suggesting multiple independent reversions from distyly back to SHD. However, this transition is not associated with the loss of ancillary polymorphisms, as all the species examined in this study well retain dimorphic traits for other floral organs such as pollens and stigmas. Moreover, most species maintain strict heteromorphic self-incompatibility, while *J. officinale* has lost or at least partially lost self-incompatibility, suggesting that distyly is not always linked to self-incompatibility.

**Conclusions:** In *Jasminum*, breakdown of distyly resulted in evolutionary transitions to stigma-height dimorphism for multiple times, suggesting that distyly is not a stable floral polymorphism under certain selective forces. These findings advance our understanding on the evolution of distyly and plant reproductive systems.

## Introduction

In angiosperm, more than half of species have evolved various outcrossing mechanisms which promote genetic exchange among individuals. Distyly is one of such powerful mechanisms facilitating outcrossing through different but reciprocal positioning of stigmas and anthers between floral morphs within a species (Sanchez et al., 2010; Costa et al., 2017; Barrett, 2019). In distylous species, the short-styled individuals (S-morph) possess high-positioned anthers and meanwhile short styles (lower stigmas), whereas the long-styled individuals (L-morph) possess low-positioned anthers and long styles (higher stigmas).

Besides the reciprocal positioning of stigmas and anthers, distylous species usually exhibit several dimorphic traits of floral organs, including pollen size and exine sculpture, stigma length, and stigma papilla length, so-called ancillary polymorphisms (Barrett, 1992; Costa et al., 2017). For example, in Primulb the S-morph flowers have rougher stigmas, and produce larger but fewer pollen grains than L-morph flowers (Darwin & Darwin, 1877; Gilmartin, 2015); in Plumbaginaceae, the exine ornamentation of L-morph pollen is coarsely reticulate, whereas that of S-morph pollen is finely reticulate (Costa et al., 2017). Most of the distylous plants also exhibit heteromorphic incompatibility (self- and intra-morph diallelic incompatibility), where stigmas of one morph only accept pollens from the other morph but reject pollens from the same morphic flowers (Charlesworth & Charlesworth, 1979).

Heterostyly (distyly and tristyly) is widespread among angiosperms and has been documented in at least 28 families, of which most families are phylogenetically distant (Barrett et al., 2000; Naiki, 2012), indicating multiple independent evolutionary origins. Two main hypotheses have been proposed for the evolutionary origin of distyly. The selfing avoidance model (Charlesworth & Charlesworth, 1979) suggests that the ancestors of distylous plants were monomorphic. Self-incompatibility first occurred in populations and were selected because of prolonged inbreeding depression, followed by the origination of stigma-height dimorphism (SHD, a polymorphism of two mating types differing in style length but not anther height), and finally the evolution of distyly with reciprocal herkogamy. In contrast, the pollen transfer model (Lloyd & Webb, 1992) proposes that the ancestors of distylous plants exhibited self-compatible approach herkogamy and later evolved reverse herkogamy mediated by a stage of SHD, and finally possessed self-incompatibility. Although the two hypotheses differ in phenotypes of hypothetical ancestors, both models suppose that the SHD is an important intermediate stage in the evolution of distyly. This evolutionary scenario is supported by the observations in Glandora (Boraginaceae), where species present a wide variety of stylar conditions and style dimorphism is probably ancestral to distyly (Ferrero et al., 2009). However, SHD is rarely observed and is thought to be difficult to maintain (Charlesworth & Charlesworth, 1979; Lloyd & Webb, 1992).

Distyly is somewhat susceptible to break down due to environmental or ecological shifts (Barrett, 2019), particularly when specialized pollinators are absent (Yuan et al., 2017; Yuan et al., 2023). In some plant lineages, such as Rubiaceae and Erythroxylum, the breakdown of distyly results in dioecy (Muenchow & Grebus, 1989). More commonly, however, it leads to monomorphic flowers, particularly homostyly (anthers and stigmas close together within a flower) accompanied by changes in ancillary polymorphisms and the breakdown of heteromorphic incompatibility. For example, in the typically distylous genus Primula, at least 45 species have been reported to be homostylous (Yuan et al., 2017; Barrett, 2019). In Primula chungensis, some homostylous populations derived from distyly eventually differentiated into populations with either approach or reverse herkogamy (Zhou et al., 2017), partially recovering the ancestral SHD phenotype, raising the possibility of reverse evolution of heterostyly. This phenomenon has only been reported in Glandora, in which the style dimorphism of G. prostrata was thought to be a reversion from distyly (Ferrero et al., 2009); however, another potential example comes from Jasminum, a genus having numerous distylous species (e.g., J. fruticans, J. odoratissimum), as well as species with SHD (e.g., J. malabaricum) (Guitián et al., 1998; Thompson & Dommée, 2000; Olesen et al., 2003; Ganguly & Barua, 2020,2021). Nevertheless, evolutionary relationships between species with distyly and SHD remain elusive due to limited investigations on the reproductive systems across this genus and the lack of robust phylogeny for the genus. Our preliminary observations of J. nudiflorum, a species morphologically much closer to Chrysojasminum than to other Jasminum species, revealed typical distylous flowers with precise reciprocal herkogamy (Supplementary Data Fig. 1a), implying the ancient origin of distyly in Jasminum. These observations prompted a broader investigation on the variation of heterostyly in Jasminum, aiming to provide a more comprehensive understanding of the evolution of heterostylous reproductive systems.

In this study, we constructed the phylogeny of 41 Jasminum species mainly occurring in southern China, and integrated data on floral syndromes of L- and S-morphs, ancillary polymorphisms, and heteromorphic incompatibility to elucidate the evolution of distyly in Jasminum. Specifically, we aim to address the following questions: (1) What is the evolutionary trajectories of distyly in Jasminum? (2) Does the SHD originate from the breakdown of distyly, or does it reflect an ephemeral state in the evolutionary transition toward distyly? (3) How do ancillary polymorphisms differ among Jasminum? (4) If distyly breaks down to SHD, how does the intra-morph incompatibility system change? This enabled us to examine the evolution and breakdown of distyly in Jasminum, uncovering previously unreported pathways of distyly breakdown. Furthermore, analyses of ancillary polymorphisms provide additional insights into the evolutionary mechanisms underlying the origin and breakdown of distyly.

## Materials and Methods

### Plant material

The genus *Jasminum* (Oleaceae) comprises more than 200 species, most of which occur in open habitats as shrubs or straggling vines (Ganguly & Barua, 2020). They usually produce yellow, white, or pink hypocrateriform corolla. In this study, we sampled 36 species from *Jasminum* and three species from *Chrysojasminum* for the phylogenetic analysis (Supplementary Data Table S1).

Nine species were further investigated for floral morph variation using morphometric method (Supplementary Data Table S2). Voucher specimens of all collected materials were deposited in the Herbarium of South China Botanical Garden.

### DNA extraction and sequencing

Total genomic DNA was extracted from silica-dried leaf samples using a modified CTAB method (Doyle & Doyle, 1987). Genome sequencing was conducted on the Illumina NovaSeq 6000 platform (Novogene Co. Ltd., Tianjin, China), with 150 bp paired-end reads and an insert size of 350 bp.

### Plastome assembly and annotation, and phylogenetic inference

The chloroplast genome for each species/specimen was assembled from clean paired-end sequencing data using the GetOrganelle pipeline (Jin *et al*., 2020). The assemblies were visualized with Bandage (Wick *et al*., 2015) and annotated using the Plastid Genome Annotator (Qu *et al*., 2019) with the *Jasminum sambac* plastome (NC_034694) as the reference. The start/stop codons, intron/exon boundaries, and tRNA genes in the preliminary annotation were manually verified and adjusted in Geneious Prime 2020.1.2 (Biomatters Ltd., Auckland, New Zealand).

Additionally, chloroplast genome sequences of four *Jasminum* species were retrieved from GenBank, including *J. fluminense* (NC_042272), *J. polyanthum* (NC_042273), *J. sambac* (NC_034694), and *J. tortuosum* (NC_034691). *Syringa pinnatifolia* (NC_041119) and *Fraxinus hubeiensis* (MT812688) were used as outgroups (Supplementary Data Table S3).

### Phylogenetic inference and reconstruction of ancestral state

Before phylogenetic analysis, all sequences were standardized in orientation following the LSC-IRb-SSC-IRa order. A data matrix was constructed using the complete chloroplast genome sequences and aligned with MAFFT v7.525 (Katoh & Standley, 2013). Phylogenetic trees were inferred using both maximum likelihood (ML) and Bayesian inference (BI) methods. ML analysis was performed using IQ-TREE v3.0.1 (Wong *et al*., 2026) with the GTR+G substitution model, and assessed with 1000 bootstrap replicates. BI analysis was conducted using MrBayes (Ronquist *et al*., 2012) with the following settings: lset nst=6; rates=gamma; mcmcp ngen=1000000; relburnin=yes; burninfrac=0.25; printfreq=1000; samplefreq=1000; nchains=4; savebrlens=yes; other settings=default.

Based on the phylogenetic tree, we reconstructed the ancestral state using Mesquite v4.03 (Maddison & Maddison, 2026). Maximum parsimony and maximum likelihood (ML) methods were used to inference the ancestral state. In our analysis, ‘0’ denotes the distyly and ‘1’ stigma-height dimorphism.

### Divergence time estimation

Divergence times within Jasmineae were estimated using the Bayesian approach in MCMCtree forked version [4.10.9] available at https://github.com/dosreislab/mcmctree (Yang & Rannala, 2006; Rannala & Yang, 2007). Chloroplast genome sequences were aligned using MAFFT v7.525 (Katoh & Standley, 2013), and subsequently trimmed by trimal v1.5.rev0 (Capella-Gutiérrez *et al*., 2009). The trimmed sequences were converted to phylip format using MEGA12 (Kumar *et al*., 2024). Because reliable fossil evidence is lacking within Jasmineae, three secondary calibration points derived from Dupin *et al*. (2024) were employed: (1) the crown node of Jasmineae and Oleeae (73.9 Mya), (2) the crown node of Jasmineae (52.3 Mya), and (3) the crown node of the clade (Ligustrinae, (Fraxininae, Oleinae)) (53.0 Mya). Skew-normal priors were assigned to all calibration points, with the offset values corresponding to the minimum ages. The ML topology inferred by IQ-TREE was used as a fixed topology. MCMCtree analyses were conducted with the following settings: clock=3; model=4; alpha=0.5; ncatG=5; print=2; burnin=20,000; sampfreq=10; nsample=500,000; other settings=default.

### Floral morphometry

To investigate the floral morphs, mature flowers were sampled during the flowering stage. For each morph of each species, ten individuals were randomly selected, and three flowers from different inflorescence of each individual were collected. Floral organs were dissected and measured to 0.01 mm using a digital calliper. The following traits were recorded (Fig. S1B): anther height, stigma height, anther length, style length, corolla tube diameter, corolla tube length, and corolla diameter.

Traits were compared between morphs using independent-samples t-tests. In order to test the relationship among these traits, Pearson correlation coefficients (*r*) among flower traits were calculated using cor function in R version 4.2.2. The overall Reciprocity Index (RI), upper sexual organs RI (the reciprocity relationships between stigmas of L-morphs and anthers of S-morphs) and lower sexual organs RI (the reciprocity relationships between anthers of L-morphs and stigmas of S-morphs) of each examined species were calculated according to Sanchez *et al*. (2008) by using the Excel macro “Recipro-v20”.

### Ancillary polymorphisms

For each morph of examined species, anthers of randomly selected flowers at full anthesis were collected and preserved in 70% formalin acetic alcohol (FAA). Anthers were broken using a tweezer to release pollens. Pollens were fixed in a mixed solution of 3% glutaraldehyde and 2% paraformaldehyde, rinsed with 0.1 M phosphate buffer, and dehydrated through a graded ethanol series (30–100%). After critical point drying using a LEICA EM CPD 300, samples were sputter-coated with approximately 10 nm of platinum (Pt) using a JFC-1600 ion sputter coater. Thirty pollen grains of each morph were randomly selected and examined using a JSM-6360LV scanning electron microscope. Images of both equatorial and polar views were obtained. The length of equatorial axis (E), polar axis (P), and area of lumina were measured by Image-J. In this study, pollen reticulations with diameters smaller than 2 μm were classified as fine reticulations, whereas those with diameters larger than 2 μm were classified as coarse reticulations.

Samples of stigmas with papillae for each floral morph were prepared and examined using the same method as described above.

### Heteromorphic incompatibility

Artificial pollination treatments were conducted according to the combinations presented in Supplementary Data Table S4.

Flowers were collected 4 or 8 h after pollination and fixed in 75% FAA. Fixed styles were softened in 10% sodium sulfite solution in a boiling water bath for 4–5 h, and subsequently stained with 0.1% aniline blue. The growth of pollen tubes was observed under a fluorescence microscope (BX41, Olympus) and was recorded using Image-Pro Plus.

## Results

### Variation of floral morphs among *Jasminum* species

To investigate the evolutionary dynamics of distyly across *Jasminum*, floral morphs of nine species across the phylogeny were examined, including *C. subhumile*, *C. humile* var. *microphyllum*, *J. nudiflorum*, *J. officinale*, *J. lanceolaria, J. beesianum, J. pentaneurum, J. tonkinense*, and *J. yuanjiangense*. All the examined species exhibited spatial separation between anthers and stigmas, indicating that herkogamy is likely well retained in *Jasminum* and *Chrysojasminum* (Fig. 1). The two *Chrysojasminum* species and four *Jasminum* species (*J. nudiflorum*, *J. officinale*, *J. beesianum* and *J. yuanjiangense*) showed more precise reciprocal herkogamy (RI > 0.7) between L- and S-morph flowers than others (Fig. 2A, Supplementary Data Fig. 2C-E; Supplementary Data Table S5). Three species, *J. lanceolaria* (RI = 0.529), *J. pentaneurum* (RI = 0.421) and *J. tonkinense* (RI = 0.026), exhibit much lower style-stamen reciprocity than typical distylous species (Fig. 2B, S2F-G; Supplementary Data Table S5). Within each of these three species, stigma heights are significantly different between floral morphs (*p* < 0.01), whereas differences in anther heights between floral morphs were much less remarkable than in typical distylous species (Supplementary Data Table S6 and S7), suggesting that they are in condition of relaxed stigma-height dimorphism in which the two floral morphs vary in style length but have similar anther height (Ferrero *et al*., 2011). Therefore, types of style polymorphism for the three species were classified as relaxed stigma-height dimorphism (hereafter also referred as stigma-height dimorphism, SHD).

**Figure 1.**
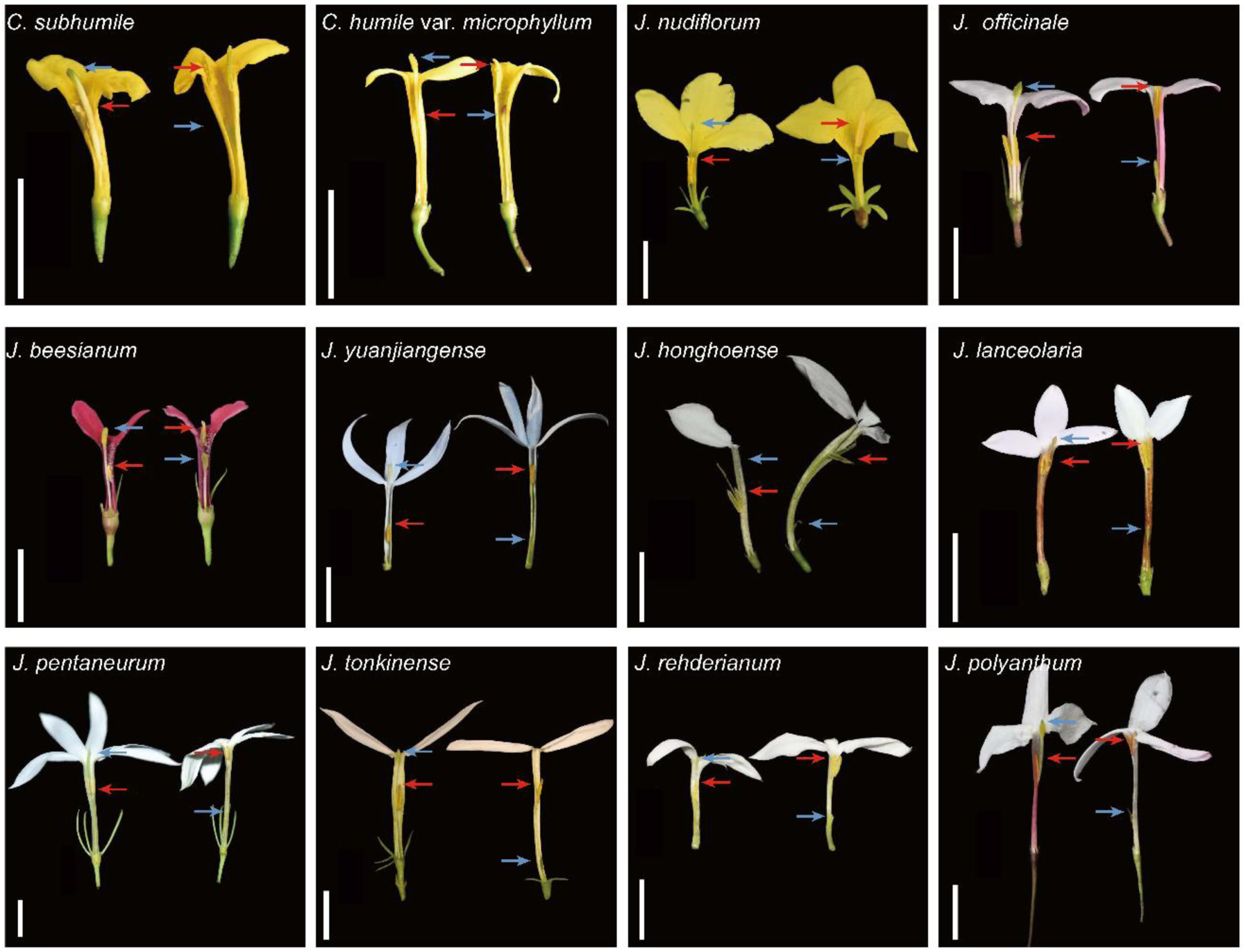
Floral morphs of Jasminum species. Chrysojasminum subhumile, C. humile var. microphyllum, Jasminum nudiflorum, J. officinale, J. beesianum, J. yuanjiangense and J. honghoense exhibited reciprocal herkogamy. Jasminum lanceolaria, J. pentaneurum, J. tonkinense, J. rehderianum, J. polyanthum exhibited reduced reciprocal herkogamy. Scale bar = 1 cm.

**Figure 2.**
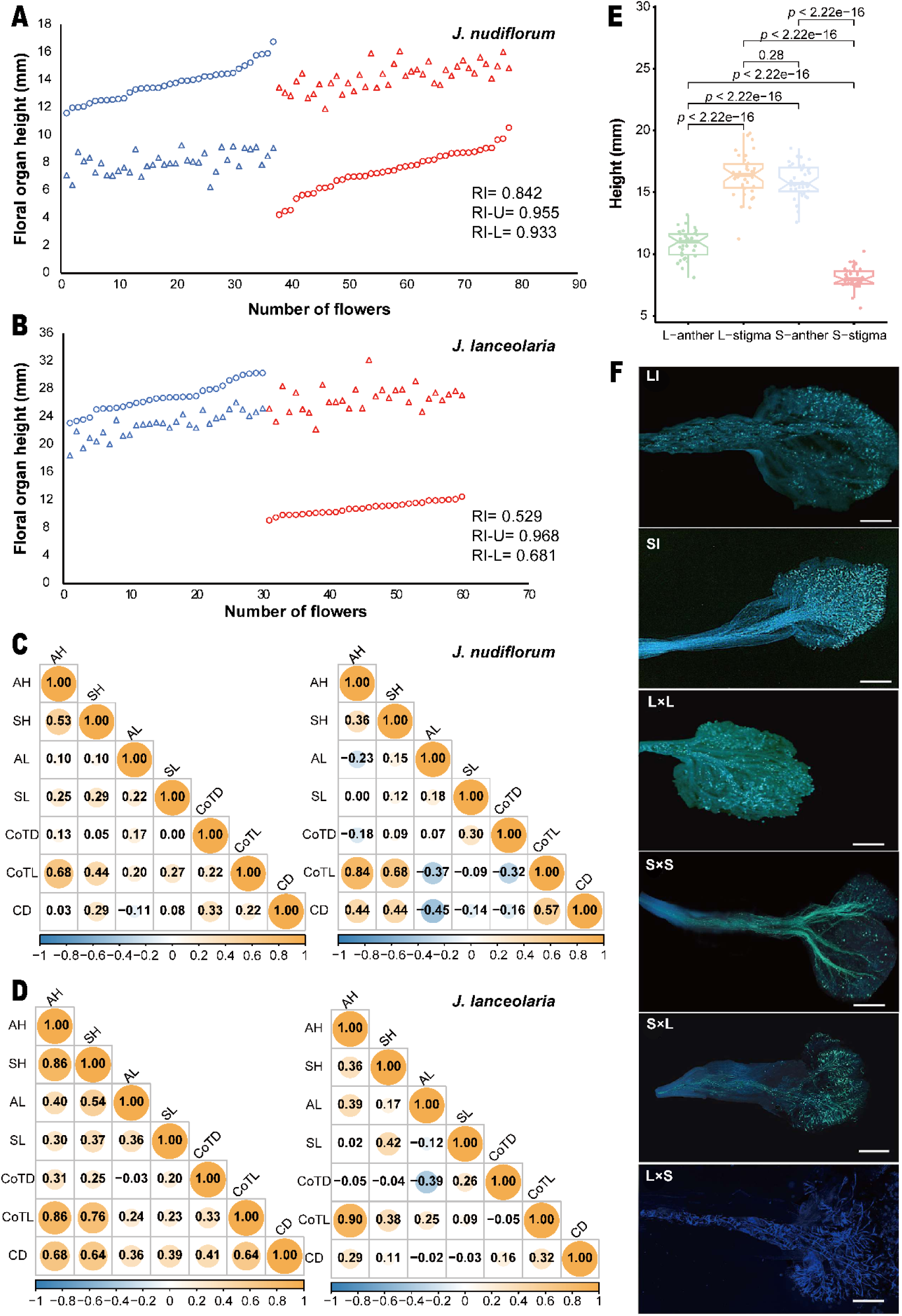
Floral trait variation, morphological correlation and self-incompatibility investigation in distylous *Jasminum* species and species with stigma-height dimorphism. (A-B) The reciprocity of *J. nudiflorum* (A) and *J. lanceolaria* (B) examined in this study. All samples are ordered by stigma height, with corresponding floral organ heights displayed. The blue and red triangles indicate anther heights of L- and S-morphs; the blue and red circles indicate stigma heights of L- and S-morphs. (C-D) Pearson correlation analysis of floral traits between L- and S-morphs for *J. nudiflorum* (C) and *J. lanceolaria* (D). Left: Flowers of L-morphs; Right: Flowers of S-morphs; CoTD: corola tube diameter; CoTL: corola tube length; CD: corolla diameter; SH: stigma height; AH: anther height; SL: stigma length; AL: anther length. (E) Boxplot comparison between anther and stigma heights for *J. officinale*. L-anther: anthers of L-morphs; L-style: stigmas of L-morphs; S-anther: anthers of S-morphs; S-style: stigmas of S-morphs. All pairwise comparisons showed extremely significant differences (*p* < 2.22 × 10⁻¹⁶), except for the comparison between L-style and S-anther. (F) Growth of pollen tubes in styles of *J. officinale* after artificial pollinations. LI: Self-pollination of L-morph flowers; L×L: Intramorph cross-pollination of L-morph flowers; SI: Self-pollination of S-morph flowers; S×S: Intramorph cross-pollination of S-morph flowers; S×L: Intermorph pollen cross-pollination treatment between L-morph flowers; L×S: Intermorph pollen cross-pollination treatment between S-morph flowers. Scale bar = 1 mm

In species which have lost precise reciprocal herkogamy, anthers in L-morph flowers were approximately twice as high as the stigmas of S-morphs and were much closer in height to anthers in relative to stigmas of S-morph flowers, although they were generally positioned at lower-level than the anthers in S-morph flowers (Supplementary Data Table S6 and S7). These results suggest that the changes of lower organs, particularly the elevation of anther heights in L-morph flowers, rather than upper organs caused the breakdown of precise reciprocal reciprocity.

We also investigated the relationship between flower size and the heights of anthers and stigmas. Anther heights in both morphs were significantly correlated with corolla tube length (average Pearson correlation coefficient *r* = 0.68 ± 0.15 for L-morph and *r* = 0.77 ± 0.16 for S-morph) (Fig. 2C,D; Supplementary Data Fig. S3). Stigma heights also correlated with corolla tube lengths in L-morph flowers (average Pearson correlation coefficient *r* = 0.53 ± 0.27), whereas this correlation was weaker in S-morphs (average *r* = 0.40 ± 0.20) (Fig. 2 C,D; Fig. S3).

Five additional species, *J. rehderianum*, *J. polyanthum*, *J. honghoense*, *J. grandiflorum*, and *J. sambac*, were also qualitatively assessed for their floral morphs in this study. Due to the limitation of the population size for these species, sufficient samples were not available for detailed morphometric analysis. Nevertheless, based on our observations, *J. honghoense* and *J. grandiflorum* were considered as distylous species; *J. rehderianum*, *J. polyanthum*, and cultivated *J. sambac* were identified as stigma height dimorphic species (Fig. 1).

### Investigation of ancillary polymorphisms

To assess the evolutionary dynamics of distyly in *Jasminum*, nine species were examined for their ancillary polymorphisms, including corolla tube lengths and diameters, lengths of anthers and stigmas. Results showed that the corolla tubes of S-morph flowers were significantly longer than those of L-morph flowers in all investigated species (Supplementary Data table S8), and most species exhibited significant difference in corolla tube diameter between the two floral morphs (Supplementary Data table S9). S-morph flowers of *J. nudiflorum*, *J. officinale*, *J. tonkinense*, and *J. yuanjiangense* possessed significantly longer anthers than L-morph flowers, while no significant differences were found in other three *Jasminum* species or in two *Chrysojasminum* species (Supplementary Data table S10). Most species showed significant differences in stigma morphology between the two floral morphs, either in stigma length or shape (Supplementary Data Fig. S4; Supplementary Data table S11). For example, in *J. nudiflorum*, S-morph flowers had spherical stigmas, while the L-morph flowers had triangular pyramid-shaped stigmas.

Six species were further examined for their pollen sizes, stigma papilla shapes, and pollen exine ornamentation. No significant differences in stigma papilla characteristics were detected based on our observations. All species produced larger pollen grains in S-morph flowers (P × E ∼45μm × 45μm) than in L-morph flowers (P × E ∼35μm × 35μm) (Supplementary Data table S12 and S13). Apparent differences in the pollen exine ornamentation were observed in four species (Fig. S4). For example, in *J. yuanjiangense*, pollen grains of S-morph flowers exhibited coarse exine ornamentation, whereas those of L-morph showed fine exine ornamentation (Supplementary Data table S14). Interestingly, this trait did not exhibit dimorphism in the typical distylous *J. nudiflorum*, suggesting that the exine ornamentation of pollen is not always associated with distylous floral polymorphism.

### Heteromorphic self-incompatibility

Heteromorphic self-incompatibility was investigated in four species, *J. yuanjiangense*, *J. nudiflorum*, *J. lanceolaria*, and *J. officinale*, through artificial pollination. Results showed that both distylous species (e.g. *J. yuanjiangense*) and stigma-height dimorphic species (e.g. *J. tonkinense*) exhibited strict self-incompatibility (Supplementary Data Fig. S5A, B). L-morph flowers of *J. nudiflorum* showed strict self- and intra-morph incompatibility, as the growth of pollen tube stopped at the stigma surface (Fig. S5C). The S-morph flowers also showed strict self-incompatibility, however, under intra-morph pollination, some pollen tubes were able to grow into the style, although eventually arrested in the middle of style, suggesting that the intra-morph incompatibility has been weakened. It is noteworthy that the self- and intra-morph incompatibility has strikingly broken down for both floral morphs of *J. officinale*, despite it is typical distyly, as pollen tubes of both self- and intra-morph pollinations grew normally in the styles (Fig. 2E, F).

### Phylogenetic relationships and divergence time estimation in *Jasminum*

In order to access the evolutionary relationships between distyly and SHD, we constructed the phylogeny of *Jasminum* species which mainly occurred in China. Forty chloroplast genomes were newly sequenced and assembled, including 37 species from *Jasminum*, and three from *Chrysojasminum* (Table S1). Phylogenetic analysis showed that the three species of *Chrysojasminum* formed a single clade which is sister to the outgroup in the phylogenetic tree, and all the *Jasminum* species formed a monophyletic lineage. *Jasminum nudiflorum*, a distylous species producing yellow flowers like those of *Chrysojasminum*, was placed at the base of *Jasminum* clade, and formed a sister lineage to all the other *Jasminum* species, indicating an early divergent lineage. The remaining *Jasminum* species were grouped into several subclades, with *J. attenuatum*, *J. officinale*, and *J. grandiflorum* forming the first diverging lineage, suggesting independent divergence and speciation events within the genus (Fig. 3). Molecular dating suggested that the lineage leading to *Jasminum* and *Chrysojasminum* diverged from the outgroup approximately 85.22 million years ago (mya). The three species of *Chrysojasminum* diversified about 13.55 mya. *Jasminum* separated from *Chrysojasminum* about 60.71 mya and diversified much earlier than the three species of *Chrysojasminum*, with *J. nudiflorum* splitting first from the other *Jasminum* species at about 48.41 mya. *J. elongatum* and *J. guangxiense* represented the most recently diverged species in our dataset, with an estimated divergence time of ∼0.17 mya (Supplementary Data Fig. S6).

**Figure 3.**
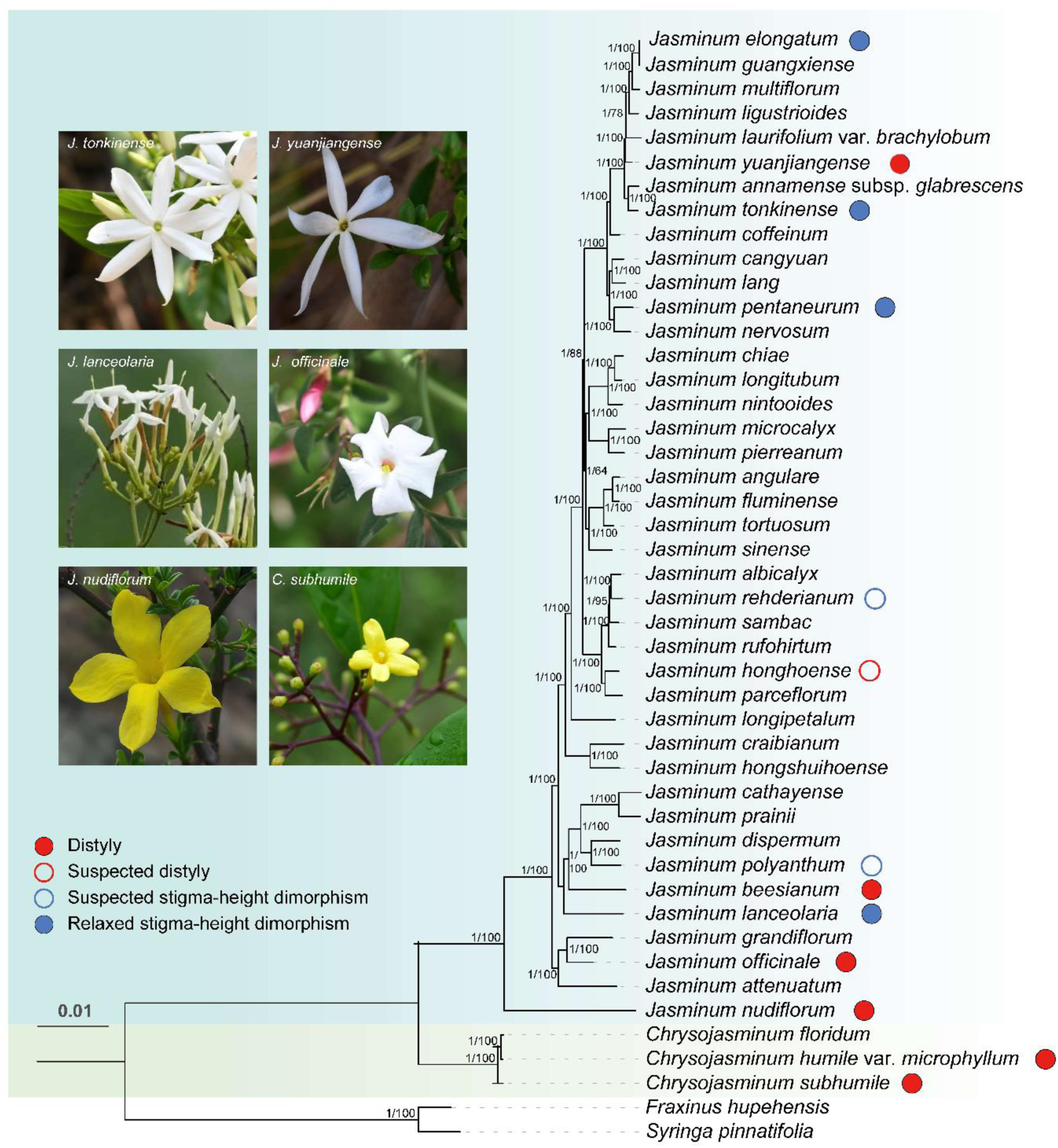
Phylogenetic relationships inferred using maximum likelihood (ML) and Bayesian inference (BI) methods. Node values indicate ML bootstrap supports and Bayesian posterior probabilities.

Ancestral reconstruction of floral polymorphism using ML methods cannot determine the ancestral state of floral traits in *Jasminum* and *Chrysojasminum* because distyly and SHD have a similar probability at most lineages (Supplementary Data Fig. S7), which is most probably due to the limited sample size in this study. However, reconstruction using parsimony methods suggest that the lineage leading to *Jasminum* and *Chrysojasminum* may have possess distylous floral polymorphism (Fig. S7). Considering that most of distylous species are located at the base of the phylogenetic tree, it is more probable that distyly has occurred in the common ancestor of *Jasminum* and *Chrysojasminum*.

## Discussion

In this study, we examined floral morphs within the tribe Jasmineae and investigated the evolutionary sequence of distyly and stigma height dimorphism through phylogenetic analysis. Our results suggested that *Jasminum* possesses both distyly and stigma-height dimorphism, and in most case exhibits distylous ancillary polymorphisms. Although we cannot determine the ancestral state using ancestral reconstruction, we suppose that distyly is the ancestral floral phenotype in the common ancestor of *Jasminum* and *Chrysojasminum*. This speculation is supported by the observation that distyly has been possessed by *Forsythia suspensa* (Ruan, 2008), a species of *Forsythia* which is out of *Jasmineae* and represents an early diverging lineage in Oleaceae (Dupin, 2024). Moreover, genomic evidences suggest that the supergene governing distyly originated in the common ancestor of *Jasminum* and *Chrysojasminum* (Raimondeau *et al*., 2024), providing strong support for our hypothesis. Therefore, in *Jasminum* it seems that SHD is derived from distyly, and distyly has independently broken down into SHD multiple times. Similar reversion is also appeared in *Glandora prostrata*, in which style dimorphism is derived from a distylous ancestor (Ferrero *et al*., 2009). In contrast, the breakdown of distyly in other plant linages most commonly results in homostyly (Mora-Carrera *et al*., 2023). Although SHD is usually considered a transitional stage in the evolution of distyly, in some plant lineages such as *Narcissus*, it can be stably maintained under certain selective mechanisms (Baker *et al*., 2000a; Zakkoumi *et al*., 2024). Collectively, these findings suggest that SHD can persist as a stable reproductive strategy that confers selective advantages in particular ecological conditions. Our results further showed that the key ancillary polymorphisms did not exhibit significant variation under the breakdown of distyly. In either distylous or stigma-height dimorphic species, pollen grains were significantly larger in S-morph flowers than in L-morph flowers. Stigmas exhibited significant dimorphism in *J. lanceolaria*, also in *J. elongatum* (Zhang *et al*., 2026), which is consistent with that observed in distylous species, such as *J. officinale* and *J. yuanjiangense*.

For all species examined in this study, anther heights were significantly correlated with corolla tube lengths. One possible explanation for this phenomenon is that the elongation of the corolla tube would increase anther heights, because the filaments are attached to the inner wall of the corolla tube. Generally, species with SHD produce longer corolla tubes than distylous species, and their anthers of L-morph flowers are much closer in height to the anthers rather than stigmas of S-morph flowers. Therefore, elongation of the corolla tube and particularly elevation of anthers in L-morphs, rather than in S-morphs, likely contributed to the loss of precise reciprocity between stigmas and anthers. This finding is supported by observations in other species, such as *J. odoratissimum* (Olesen *et al*., 2003). Changes in L-morph anther position may be of functional significance for pollination and mating. Considering that most of stigma-height dimorphic species commonly possess extremely longer corolla tubes, we hypothesize that the lower anther position may impede pollinators to pick up pollens and further cause pollen loss when pollinators remove pollens out of corolla tubes; elevated anther heights near the top of corolla tube could alleviate this obstacle and therefore improve pollen transfer. However, the exact ecological factors and fitness advantages associated with these phenotypes remain to be explored.

Recent studies have uncovered the genetic mechanisms underlying distyly in several plant lineages (Luo *et al*., 2025). Mutations in *S*-locus-linked genes controlling either style length or anther position have been shown to cause elongation of styles (long homostyles) or loss of anther elevation (short homostyles) in S-morphs (Huu *et al*., 2016; Zhao *et al*., 2023). In *Chrysojasminum fruticans* (Jasmineae), there is evidence that the distyly is also controlled by an S-morph-dominant hemizygous supergene (Raimondeau *et al*., 2024). Therefore, if the distyly in *Jasminum* is governed by the same genetic mechanism, phenotypic changes observed in L-morph flowers are unlikely to be caused by mutations of the *S*-locus supergene, which should be absent from L-morphs according to the hemizygous model.

It has been consistently revealed that the *S*-locus supergene is tightly involved in the control of either female or male self-incompatibility in S-morph flowers (Matsui & Yasui, 2020; Matzke *et al*., 2021; Huu *et al*., 2022; Gutiérrez-Valencia *et al*., 2024). Therefore, breakdown of heterostyly is often accompanied by the loss of self-incompatibility (Wang *et al*., 2020). In contrast, our result showed that breakdown of structural integrity did not result in the loss of heteromorphic incompatibility. For example, *J. elongatum*, a species with SHD, still exhibit strong self- or intra-morph incompatibility (Zhang *et al*., 2026). Interestingly, *J. officinale*, a distylous species with precise reciprocal herkogamy, has lost self- or intra-morph incompatibility. Similarly, some typical distylous species, e.g., *Primula oreodoxa* and *Cordia subcordata*, have lost heteromorphic incompatibility while retaining typical distylous floral syndrome (Yuan et al., 2019; Shi *et al*., 2026). These observations suggest that self-incompatibility is not always required for the persistence of heterostyly. Indeed, a simulation in *Narcissus*, a genus dimorphic for stigma height without heteromorphic incompatibility, suggests that the SHD can be steadily maintained when levels of disassortative mating are greater than assortative mating (Baker *et al*., 2000b). It is likely that other genes out of *S*-locus supergene may participate in controlling heteromorphic incompatibility, and mutations occurred in these genes may have altered the incompatibility in these species (Shi *et al*., 2026). Nevertheless, further works involving genome sequencing and comparative genomic analysis are needed to provide valuable information for understanding the molecular mechanisms underlying the breakdown of either floral morphology or heteromorphic incompatibility.

## Conflicts of Interest

The authors declare no conflict of interest relevant to the publication of this work.

## Author contributions

DZ and ZZ designed and supervised this study; SC, KZ, and JZ performed field investigations, data collections, and phylogenetic analyses; SC, KZ, and XP contributed to the morphological studies; YZ, WX, SL, ZM and TT participated in data analysis; SC, ZZ and MS prepared the draft; ZZ, DZ, and MS revised the manuscript.

## Availability of Data and Materials

All the data that support the findings of this study have been included in Figs S1–S7, and Tables S1–S14.

## Acknowledgements

This work was supported by grants from the Southwest and Beibu Gulf Scientific Expedition Phase I Program (Grant No. 2026XNKK00124), the Guangdong Provincial Special Fund for Natural Resource Affairs on Ecology and Forestry Construction (GDZZDC20228704), and grants from Guangxi (Bagui) Outstanding Young Talents Cultivation Program.

## Notes

### Competing Interest Statement

The authors have declared no competing interest.

