## Supplementary figures and images for "From distyly to stigma-height dimorphism: phylogenetic insights into the evolutionary breakdown of distyly in *Jasminum*"

### Supplemental Figure S3

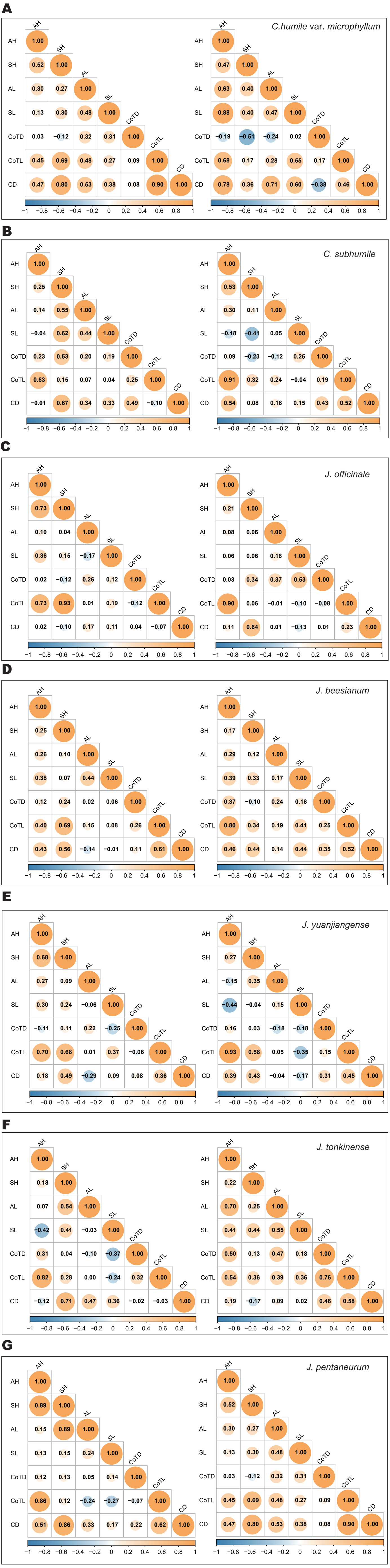
