## Supplemental tables and figures for "From distyly to stigma-height dimorphism: phylogenetic insights into the evolutionary breakdown of distyly in *Jasminum*"

The following Supporting Information is available for this article:

**Fig. S1** Longitudinal anatomical diagrams of flowers of seven species, showing the reciprocal positioning of anthers and stigmas between two floral morphs.

**Fig. S2** The reciprocity of *Jasminum* species examined in this study. All samples are ordered by stigma height, with corresponding floral organ heights displayed.

**Fig. S3** Pearson correlation analysis of floral traits within floral morphs.

**Fig. S4** Micrographs for pollen grains and stigmas from L- and S-morph flowers in *Jasminum* species.

**Fig. S5** Growth of pollen tubes after artificial pollination.

**Fig. S6** The phylogenic tree with divergence time showing the relationships among species and their estimated divergence times.

**Fig. S7** Inference of ancestral sexual polymorphisms in *Jasminum*.

**Table S1** Summary for the taxa sampled for the phylogenetic analysis in this study.

**Table S2** Jasminum species used for floral morphometry in the present study.

**Table S3** Accession numbers for chloroplast genomes downloaded from GenBank.

**Table S4** Summary of artificial pollination.

**Table S5** The reciprocity indices between the L-morph and S-morph.

**Table S6** The comparison of Anther heights between L-Morph and S-Morph.

**Table S7** The comparison of Stigma heights between L-Morph and S-Morph.

**Table S8** The comparison of Corolla tube lengths between L-Morph and S-Morph.

**Table S9** The comparison of Corolla tube diameters between L-Morph and S-Morph.

**Table S10** The comparison of Anther lengths between L-Morph and S-Morph.

**Table S11** The comparison of Style lengths between L-Morph and S-Morph.

**Table S12** The comparison of Polar axes between pollen from L-Morph and S-Morph.

**Table S13** The comparison of Equatorial axes of pollen between L-Morph and S-Morph.

**Table S14** The comparison of Lumen areas between pollen from L‑morph and S‑morph.

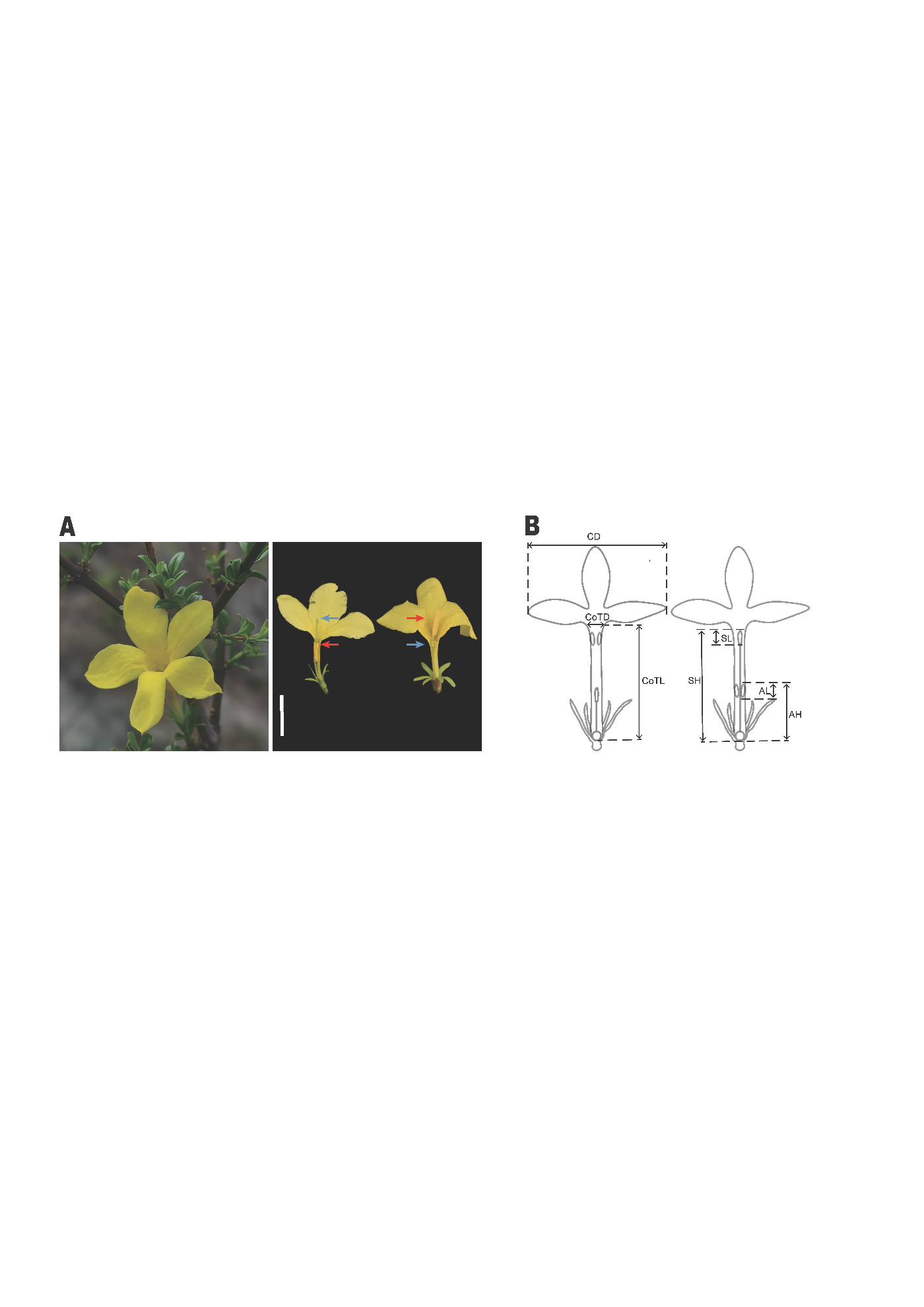

**Fig. S1** Floral morphs of *Jasminum* species. (A) Anatomical illustration of L- and S-morph flowers in *J. nudiflorum* with precise reciprocal herkogamy. (B) Floral measurements of L- and S-morphs illustrating the key floral organ dimensions. CoTD: corola tube diameter; CoTL: corola tube length; CD: corolla diameter; SH: stigma height; AH: anther height; SL: stigma length; AL: anther length. Scale bar = 1 cm.

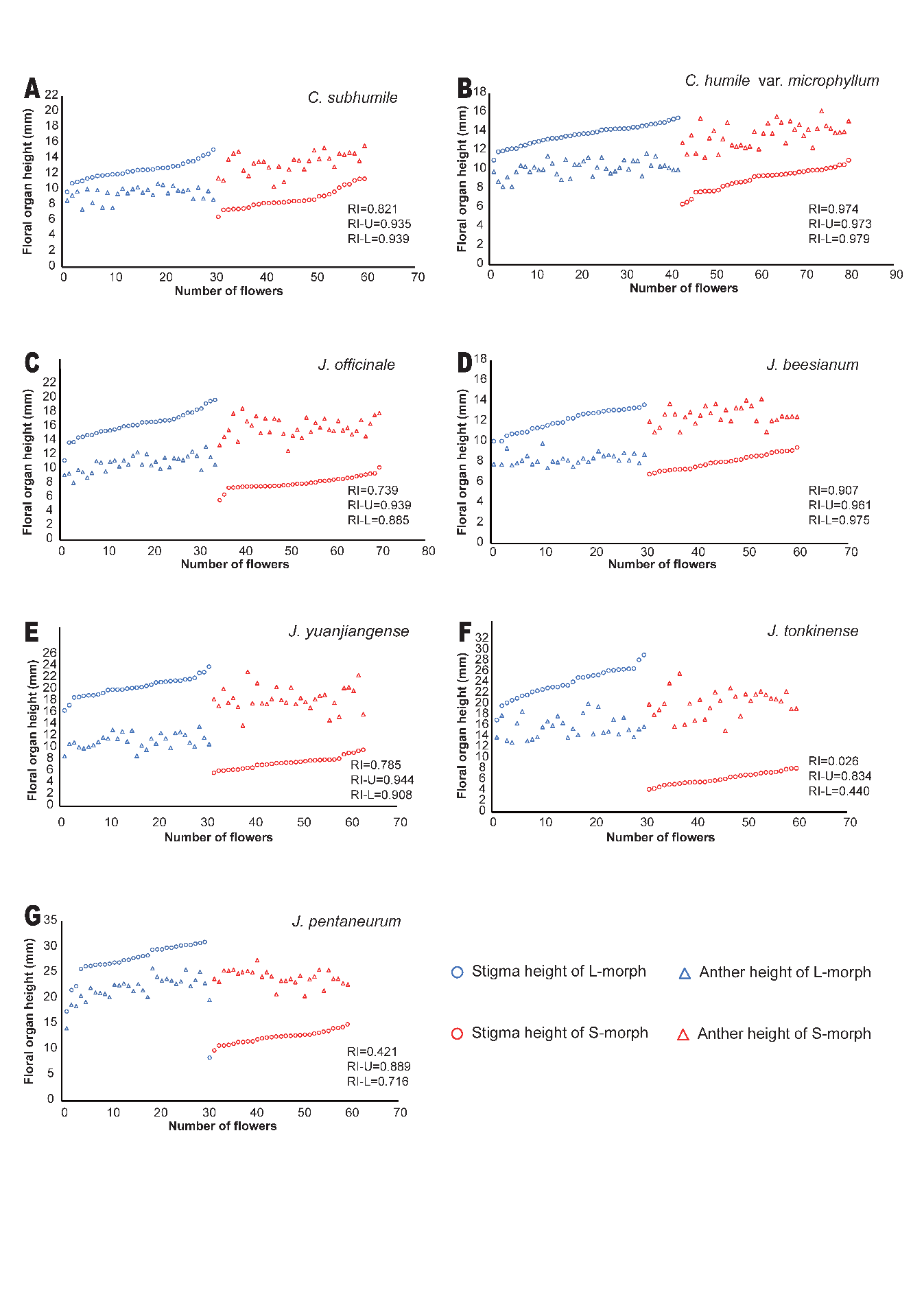

**Fig. S2** The reciprocity of *Jasminum* species examined in this study. All samples are ordered by stigma height, with corresponding floral organ heights displayed. The blue and red triangles indicate anthers height of L- and S-morph; the blue and red circles indicate stigmas height of L- and S-morph. RI: the reciprocity index of total stigmas and anthers; RI-U: the reciprocity index of upper organs (the stigmas of L-morphs and the anthers of S-morphs); RI-L: the reciprocity index of lower organs (the stigmas of S-morphs and the anthers of L-morphs)

**Fig. S3** Pearson correlation analysis of floral traits within floral morphs. Left: Flowers of L-morph; Right: Flowers of S-morph; CoTD: corolla tube diameter; CoTL: corolla tube length; CD: corolla diameter; SH: stigma height; AH: anther height; SL: stigma length; AL: anther length.

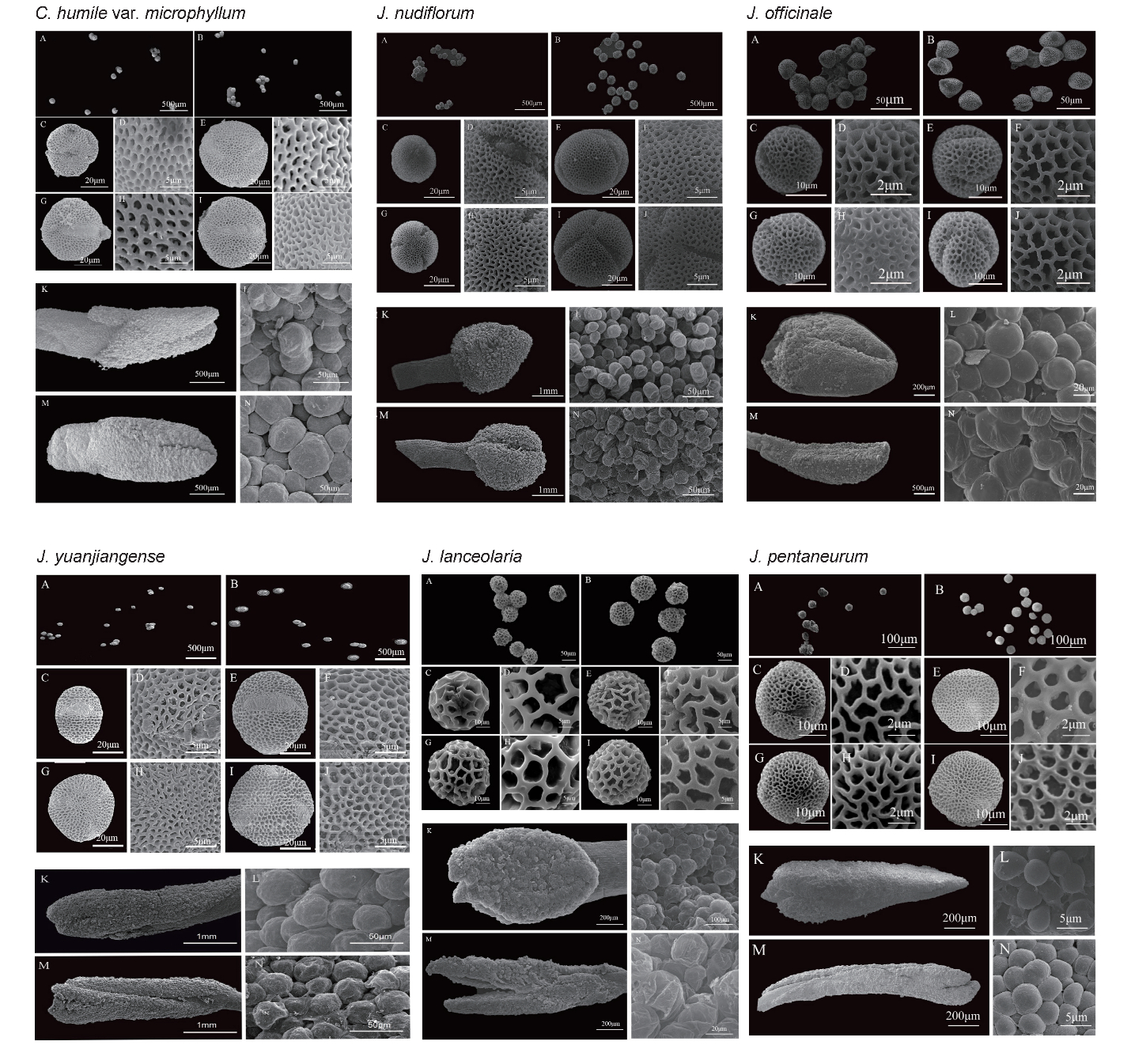

**Fig. S4** Micrographs for pollen grains and stigmas from L- and S-morph flowers in *Jasminum* species. For each species: (A, B) Pollen grains of L- and S-morph flowers; (C, D) Equatorial views of pollen grains in L-morph flowers; (E, F) Equatorial views of pollen grains in S-morph flowers; (G, H) Polar views of pollen grains in L-morph flowers; (I, J) Polar views of pollen grains in S-morph flowers. (K, M) Stigma of L- and S-morph flowers; (L, N) Papillae cells of L- and S-morph flowers. Scale bar = 1 mm.

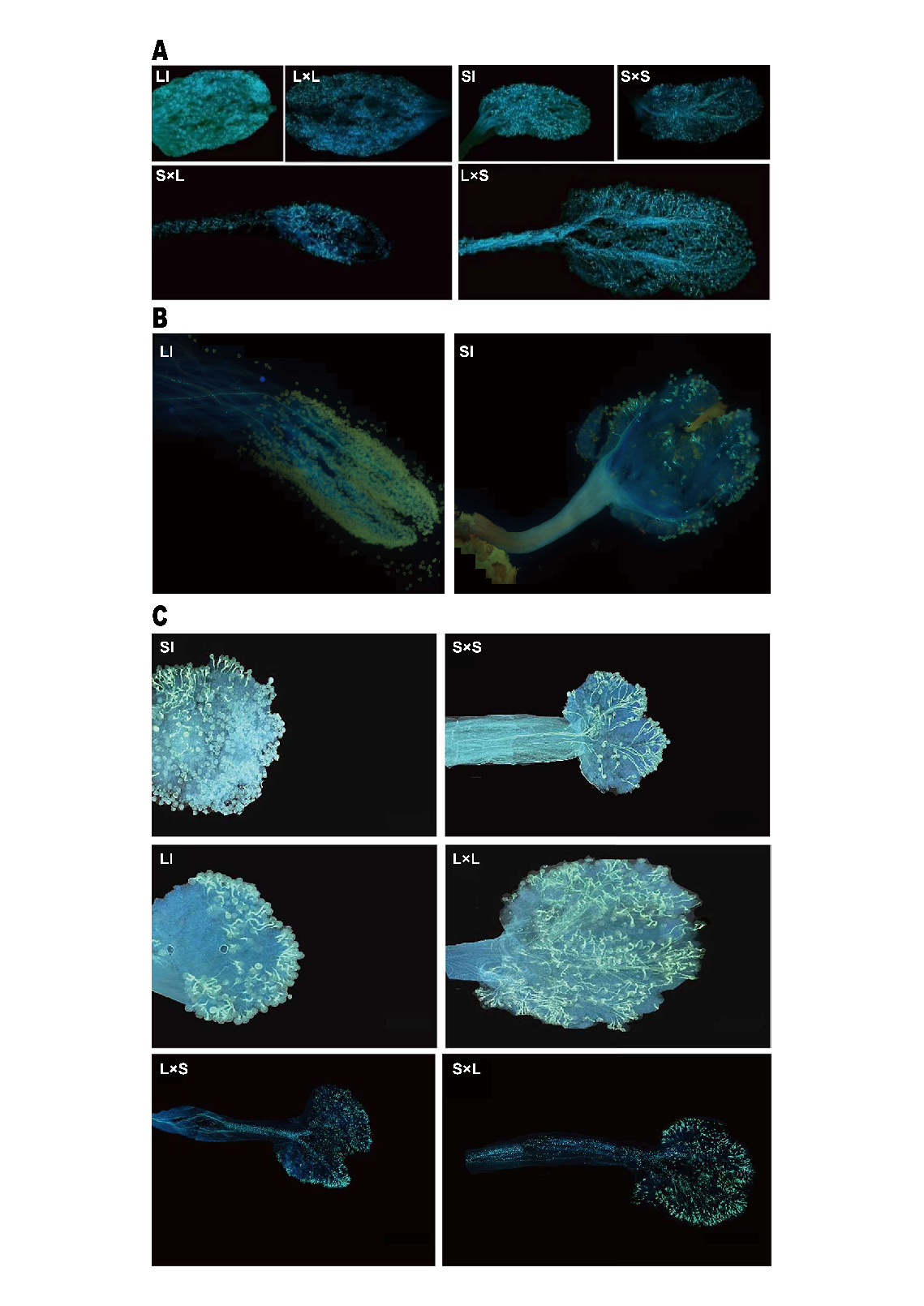

**Fig. S5** Growth of pollen tubes after artificial pollination of *J. yuanjiangense* (A), *J. tonkinense* (B) and, *J. nudiflorum* (C). LI: Self-pollination treatment in L-morph flowers; L×L: Intramorph pollen cross-pollination treatment in L-morph flowers; SI: Self-pollination treatment in S-morph flowers; S×S: Intramorph pollen cross-pollination treatment in S-morph flowers; S×L: Intermorph pollen cross-pollination treatment between L-morph flowers; L×S: Intermorph pollen cross-pollination treatment between S-morph flowers.

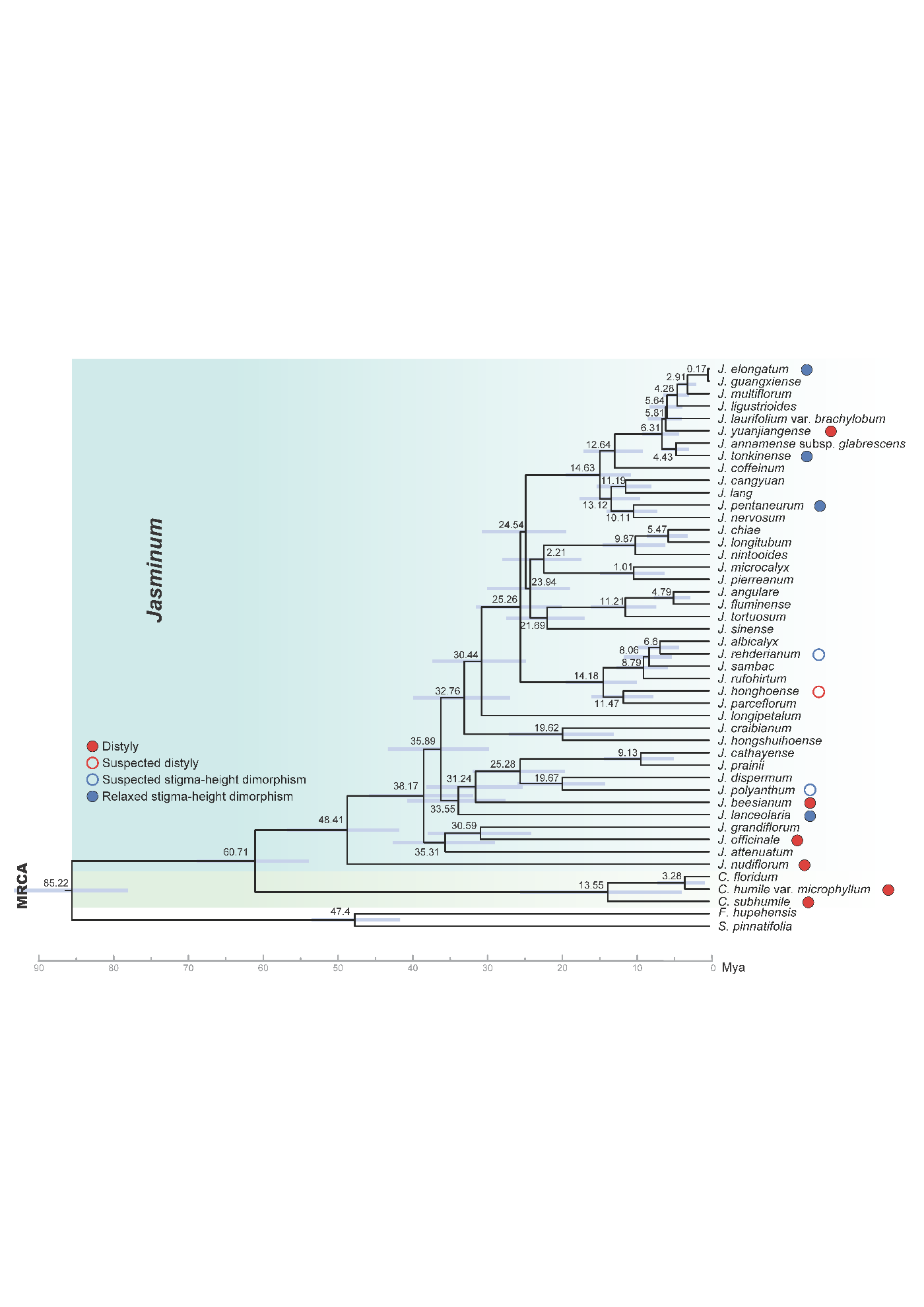

**Fig. S6** The phylogenetic tree with divergence time showing the relationships among species and their estimated divergence times. Node bars represent 95% highest posterior density (HPD) intervals, and numbers at nodes indicate divergent time.

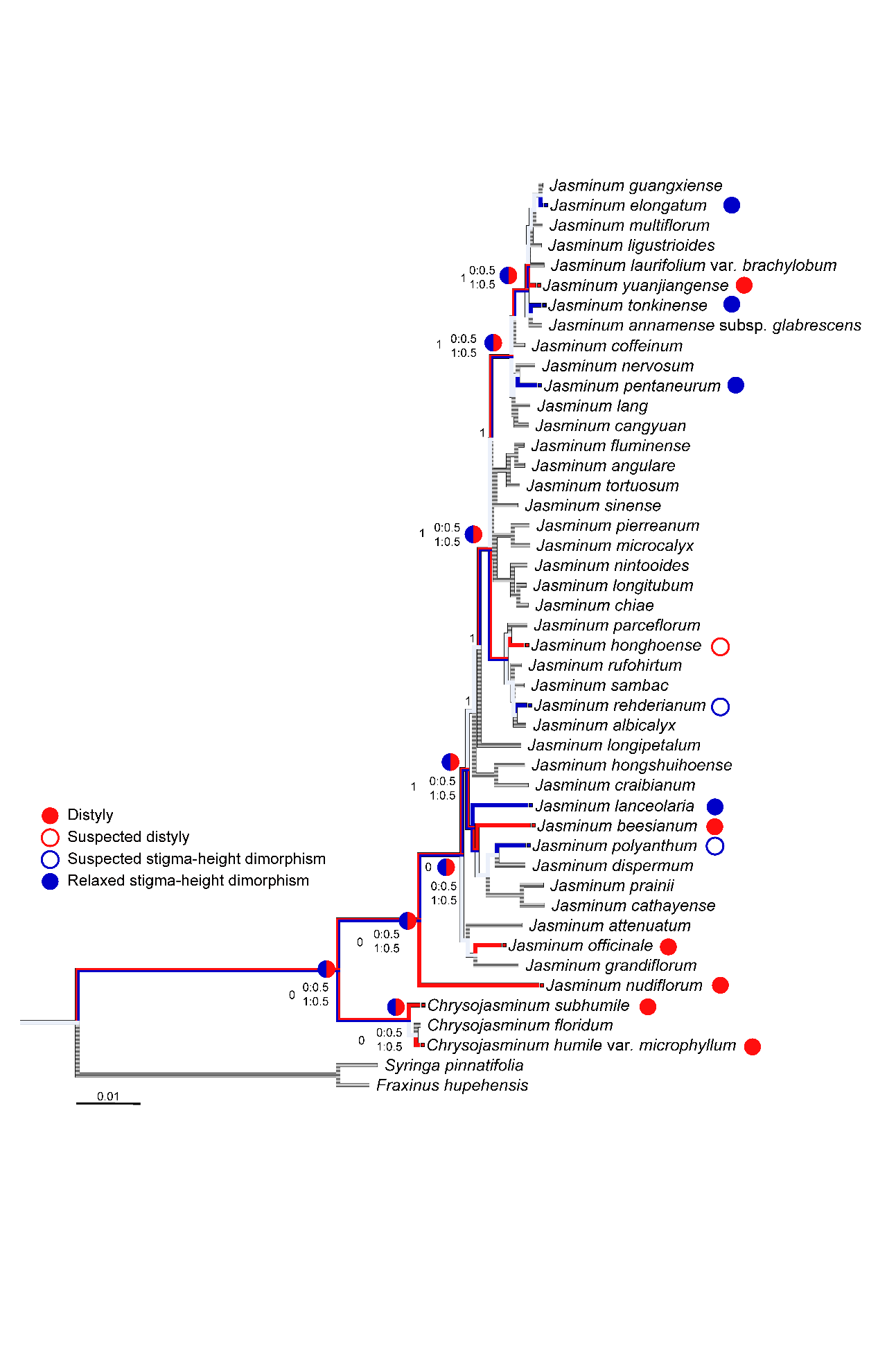

**Fig. S7** Inference of ancestral sexual polymorphisms in *Jasminum*. The most parsimonious states with probabilities of maximum likelihood method are shown at major branches. 0: distyly, 1: relaxed stigma-height dimorphism.

**Table S1 Summary for the taxa sampled for the phylogenetic analysis in this study.**

| Species | Vocher (IBSC) | Sampling location |
| --- | --- | --- |
| *Chrysojasminum subhumile* Banfi & Galasso | Kai Zhang 00164 | Yuanjiang, Yunnan, China |
| *Chrysojasminum floridum* (Bunge) Banfi | Kai Zhang 01003 | Tengzhou, Shandong, China |
| *Chrysojasminum humile* (L.) Banfi | Kai Zhang 01190 | Bomi, Xizang, China |
| *Jasminum annamense* subsp. *glabrescens* | Kai Zhang 00556 | Mengla, Yunnan, China |
| *Jasminum attenuatum* Roxb. & G. Don | Kai Zhang 01179 | Jinghong, Yunnan, China |
| *Jasminum beesianum* Forrest & Diels | Kai Zhang 01180 | Yulong, Yunnan, China |
| *Jasminum* *cangyuan* | Kai Zhang 01171 | Cangyuan, Yunnan, China |
| *Jasminum cathayense* Chun ex Chia | Kai Zhang 01209 | Yunkaishan, Guangdong, China |
| *Jasminum chiae* Kai Zhang bis & D.X. Zhang | Kai Zhang 00612 | Cangyuan, Yunnan, China |
| *Jasminum coffeinum* Hand.-Mazz. | Kai Zhang 00331 | Ningming, Guangxi, China |
| *Jasminum craibianum* Kerr | Kai Zhang 01069 | Baoting, Hainan, China |
| *Jasminum dispermum* Wall. | Youpai Zeng ZYP30 | Wuliangshan, Yunnan, China |
| *Jasminum elongatum* (Bergius) Willd. | Kai Zhang 01146 | Ningming, Guangxi, China |
| *Jasminum grandiflorum* L. | Kai Zhang 00549 | Mengla, Yunnan, China |
| *Jasminum rufohirtum* C.B. Clarke | Kai Zhang 00590 | Menglian, Yunnan, China |
| *Jasminum guangxiense* B.M. Miao | Kai Zhang 00337 | Ningming, Guangxi, China |
| *Jasminum honghoense*ng | Kai Zhang 00405 | Honghe, Yunnan, China |
| *Jasminum hongshuihoense* Jien ex B.M. Miao | Kai Zhang 01154 | Longzhou, Guangxi, China |
| *Jasminum lanceolaria* Roxb. | Kai Zhang 01206 | Tongshan, Hubei, China |
| *Jasminum lang* Gagnep. | Kai Zhang 01144 | Ningming, Guangxi, China |
| *Jasminum laurifolium* Roxb. | Kai Zhang 00639 | Cangyuan, Yunnan, China |
| *Jasminum ligustroides* Chia | Kai Zhang 01085 | Danzhou, Hainan, China |
| *Jasminum longipetalum* King & Gamble | Kai Zhang 01062 | Wanning, Hainan, China |
| *Jasminum longitubum* | Kai Zhang 00301 | Longzhou, Guangxi, China |
| *Jasminum microcalyx* Hance | Kai Zhang 01055 | Wanning, Hainan, China |
| *Jasminum multiflorum* (Burm. f.) Andrews | Kai Zhang 01008 | Guangzhou, Guangdong, China |
| *Jasminum nervosum* Lour. | Kai Zhang 01079 | Exianling, Hainan, China |
| *Jasminum nintooides* Rehder | Kai Zhang 00425 | Wenshan, Yunnan, China |
| *Jasminum nudiflorum* Lindl. | Kai Zhang 01005 | Tengzhou, Shandong, China |
| *Jasminum officinale* L. | Kai Zhang 01021 | Maerkang, Sichuan, China |
| *Jasminum parceflorum* | Kai Zhang & Mingsong Wu 00533 | Jinghong, Yunnan, China |
| *Jasminum albicalyx* | Kai Zhang 00358 | Longzhou, Guangxi, China |
| *Jasminum pentaneurum* Hand.-Mazz. | Kai Zhang 00199 | Zhanjiang, Guangdong, China |
| *Jasminum pierreanum* Gagnep. | Kai Zhang 01073 | Jianfengling, Hainan, China |
| *Jasminum prainii* H.Lév. | Kai Zhang 01186 | Xishui, Guizhou, China |
| *Jasminum rehderianum* Kobuski | Kai Zhang 01075 | Danzhou, Hainan, China |
| *Jasminum sinense* Hemsl. | Kai Zhang 01184 | Anlong, Guizhou, China |
| *Jasminum tonkinense* Gagnep. | Kai Zhang 00356 | Longzhou, Guangxi, China |
| *Jasminum yuanjiangense* P.Y. Bai | Kai Zhang 00166 | Yuanjiang, Yunnan, China |

**Table S2 Jasminum species used for floral morphometry in the present study.**

| Species | Longitude, latitude | Collection place |
| --- | --- | --- |
| *Chrysojasminum subhumile* | 23°32'15.1289"N, 102°14'29.3291"E | Yunnan Province, China |
| *Chrysojasminum humile* var. *microphyllum* | 28°28'3.1"N, 98°55'1.4"E | Yunnan Province, China |
| *Jasminum nudiflorum* | 28°37′N ，98°00′W | Xizang Autonomous Region, China |
| *Jasminum officinale* | 25°41′26″N ，100°9′45″W | Yunnan Province, China |
| *Jasminum lanceolaria* | 24°14′33″N ，112°45′13″W | Guangdong Province, China |
| *Jasminum beesianum* | 26°59′44″N ，100°11′50″E | Yunnan Province, China |
| *Jasminum pentaneurum* | 20°35′N ，109°31′W | Guangdong Province, China |
| *Jasminum tonkinense* | 23°23′1″N ，102°50′51″E | Yunnan Province, China |
| *Jasminum yuanjiangense* | 23°22'11.4"N, 102°25'34.1"W | Yunnan Province, China |

**Table S3 Accession numbers for chloroplast genomes downloaded from GenBank.**

| Species | Accession number |
| --- | --- |
| *Jasminum angulare* | OR605726.1 |
| *Jasminum fluminense* | NC_042272.1 |
| *Jasminum polyanthum* | NC_042273 |
| *Jasminum tortuosum* | NC_034691 |
| *Jasminum sambac* | NC_034694 |
| *Fraxinus hupehensis* | MT812688 |
| *Syringa pinnatifolia* | NC_041119 |

**Table S4 Summary of artificial pollination.**

| Treatment type | pollen parent | female parent | Cross type |
| --- | --- | --- | --- |
| Selfing | L-morph | L-morph | Self |
| Selfing | S-morph | S-morph | Self |
| Intra-morph | L-morph | L-morph | Intra-morph cross |
| Intra-morph | S-morph | S-morph | Intra-morph cross |
| Inter-morph | L-morph | S-morph | Inter-morph cross |
| Inter-morph | S-morph | L-morph | Inter-morph cross |

**Table S5 Reciprocity indices between the L-morph and S-morph.**

| Species | RI | RI-U | RI-L |
| --- | --- | --- | --- |
| *Chrysojasminum subhumile* | 0.821 | 0.935 | 0.939 |
| *Chrysojasminum humile* var. *microphyllum* | 0.974 | 0.973 | 0.979 |
| *Jasminum nudiflorum* | 0.842 | 0.955 | 0.933 |
| *Jasminum officinale* | 0.739 | 0.939 | 0.885 |
| *Jasminum lanceolaria* | 0.529 | 0.968 | 0.681 |
| *Jasminum beesianum* | 0.907 | 0.961 | 0.975 |
| *Jasminum pentaneurum* | 0.421 | 0.889 | 0.716 |
| *Jasminum tonkinense* | 0.026 | 0.834 | 0.440 |
| *Jasminum yuanjiangense* | 0.785 | 0.944 | 0.908 |

RI: the reciprocity index of total stigmas and anthers; RI-U: the reciprocity index of upper organs (the stigmas of L-morphs and the anthers of S-morphs); RI-L: the reciprocity index of lower organs (the stigmas of S-morphs and the anthers of L-morphs)

**Table S6 Comparisons of anther heights between L-morph and S-morph.**

| Species | L-Morph  X±SD (mm) | S-Morph  X±SD (mm) | t-value |
| --- | --- | --- | --- |
| *Chrysojasminum subhumile* | 9.47±0.87 | 13.46±1.34 | -13.67** |
| *Chrysojasminum humile* var. *microphyllum* | 9.46±0.73 | 13.19±0.86 | -18.04** |
| *Jasminum nudiflorum* | 7.89±0.78 | 13.97±0.95 | -27.11** |
| *Jasminum officinale* | 10.74±1.10 | 15.81±1.36 | -15.90** |
| *Jasminum lanceolaria* | 22.853±1.82 | 26.61±1.90 | -7.84** |
| *Jasminum beesianum* | 8.31±0.55 | 12.59±0.88 | -22.56** |
| *Jasminum pentaneurum* | 22.06±2.37 | 23.98±1.68 | -3.63** |
| *Jasminum tonkinense* | 15.75±1.97 | 20.16±2.47 | -7.59** |
| *Jasminum yuanjiangense* | 11.29±1.19 | 18.5±2.04 | -16.74** |

Negative t-values indicate that anther heights of S-Morph are higher than those of the L-Morph. *: *p* < 0.05, **: *p* < 0.01.

**Table S7 Comparisons of stigma heights between L-morph and S-morph.**

| Species | L-Morph X±SD (mm) | S-Morph X±SD (mm) | t-value |
| --- | --- | --- | --- |
| *Chrysojasminum subhumile* | 12.50±1.177 | 8.78±1.28 | 11.70** |
| *Chrysojasminum humile* var. *microphyllum* | 13.79±1.16 | 9.98±0.86 | 14.50** |
| *Jasminum nudiflorum* | 13.29±0.86 | 6.83±1.17 | 24.28** |
| *Jasminum officinale* | 15.95±1.5 | 7.82±0.65 | 27.26** |
| *Jasminum lanceolaria* | 26.79±2.08 | 10.87±0.89 | 38.52** |
| *Jasminum beesianum* | 12.11±1.07 | 8.05±0.76 | 16.98** |
| *Jasminum pentaneurum* | 27.68±2.99 | 12.35±1.41 | 25.35** |
| *Jasminum tonkinense* | 23.93±2.69 | 6.21±1.12 | 33.19** |
| *Jasminum yuanjiangense* | 18.8±1.46 | 7.48±0.93 | 41.35** |

Positive t-values indicate that stigma heights of L-morph are higher than those of the S-morph. *: *p* < 0.05, **: *p* < 0.01.

**Table S8 Comparisons of corolla tube lengths between L-corph and S-morph**

| Species | L-Morph X±SD (mm) | S-Morph X±SD (mm) | t-value |
| --- | --- | --- | --- |
| *Chrysojasminum subhumile* | 11.59±0.96 | 12.15±1.08 | -2.11* |
| *Chrysojasminum humile* var. *microphyllum* | 11.09±0.81 | 12.03±0.83 | -4.44** |
| *Jasminum nudiflorum* | 10.51±0.74 | 12.57±0.92 | -9.5** |
| *Jasminum officinale* | 12.76±1.91 | 14.44±1.35 | -3.92** |
| *Jasminum lanceolaria* | 24.88±1.91 | 27.31±2.04 | -4.78** |
| *Jasminum beesianum* | 10.66±0.80 | 11.53±0.94 | -3.88** |
| *Jasminum pentaneurum* | 24.88±2.17 | 25.31±3.34 | -9.03** |
| *Jasminum tonkinense* | 23.21±2.16 | 24.98±2.54 | -2.88** |
| *Jasminum yuanjiangense* | 18.84±1.29 | 19.89±0.85 | -3.72** |

Negative t-values indicate that corolla tube lengths of S-morph are longer than those of L-morph. *: *p* < 0.05, **: *p* < 0.01.

**Table S9 Comparisons of corolla tube diameters between L-morph and S-morph.**

| Species | L-Morph X±SD (mm) | S-Morph X±SD (mm) | t-value |
| --- | --- | --- | --- |
| *Chrysojasminum subhumile* | 2.26±0.32 | 1.94±0.48 | 3.06** |
| *Chrysojasminum humile* var. *microphyllum* | 1.71±0.32 | 1.87±0.31 | -2.03* |
| *Jasminum nudiflorum* | 2.43±0.17 | 2.51±0.23 | -1.53 |
| *Jasminum officinale* | 2.51±0.23 | 2.43±0.17 | 2.57* |
| *Jasminum lanceolaria* | 2.28±0.27 | 2.42±0.20 | -2.28* |
| *Jasminum beesianum* | 2.29±0.34 | 2.31±0.39 | -0.22 |
| *Jasminum pentaneurum* | 2.16±0.11 | 2.20±0.26 | -0.98 |
| *Jasminum tonkinense* | 1.81±0.22 | 2.17±0.48 | -3.63** |
| *Jasminum yuanjiangense* | 1.98±0.4 | 2.1±0.53 | -1.02 |

Positive/negative t-values indicate that corolla tube diameters of S-morph are larger/smaller than those of L-morph. *: *p* < 0.05, **: *p* < 0.01.

**Table S10 Comparisons of anther lengths between L-morph and S-morph.**

| Species | L-Morph X±SD (mm) | S-Morph X±SD (mm) | t-value |
| --- | --- | --- | --- |
| *Chrysojasminum subhumile* | 3.22±0.33 | 3.05±0.35 | 1.96 |
| *Chrysojasminum humile* var. *microphyllum* | 3.24±0.41 | 3.34±0.35 | -1 |
| *Jasminum nudiflorum* | 3.02±0.29 | 3.80±0.47 | -7.76** |
| *Jasminum officinale* | 4.63±0.22 | 5.05±0.26 | -6.64** |
| *Jasminum lanceolaria* | 4.72±0.47 | 4.64±0.45 | 0.72 |
| *Jasminum beesianum* | 2.79±0.40 | 2.61±0.38 | 1.82 |
| *Jasminum pentaneurum* | 4.48±0.48 | 4.52±0.55 | -1.03 |
| *Jasminum tonkinense* | 4.65±0.76 | 5.42±0.93 | -3.49** |
| *Jasminum yuanjiangense* | 3.21±0.35 | 3.6±0.38 | -4.13** |

Positive/negative t-values indicate that anther lengths of S-morph are longer/shorter than those of L-morph. *: *p* < 0.05, **: *p* < 0.01.

**Table S11 Comparisons of style lengths between L-morph and S-morph.**

| Species | L-Morph X±SD (mm) | S-Morph X±SD (mm) | t-value |
| --- | --- | --- | --- |
| *Chrysojasminum subhumile* | 1.66±0.34 | 1.86±0.34 | -2.23* |
| *Chrysojasminum humile* var. *microphyllum* | 1.52±0.17 | 1.60±0.29 | -1.05 |
| *Jasminum nudiflorum* | 1.11±0.21 | 1.02±0.19 | 1.65 |
| *Jasminum officinale* | 2.29±0.19 | 3.97±0.33 | -23.89** |
| *Jasminum lanceolaria* | 2.07±0.32 | 3.32±0.52 | -11.13** |
| *Jasminum beesianum* | 1.67±0.28 | 1.70±0.27 | -0.55 |
| *Jasminum pentaneurum* | 5.12±0.43 | 4.9±0.42 | 0.82 |
| *Jasminum tonkinense* | 2.39±0.80 | 2.70±0.43 | -1.84 |
| *Jasminum yuanjiangense* | 2.12±0.36 | 2.56±0.4 | -4.54** |

Positive/negative t-values indicate that style lengths of S-morph are longer/shorter than those of L-morph. *: *p* < 0.05, **: *p* < 0.01.

**Table S12 Comparisons of polar axes between pollen from L-morph and S-morph.**

| Species | L-Morph X±SD (μm) | S-Morph X±SD (μm) | t-value |
| --- | --- | --- | --- |
| *Chrysojasminum humile* var. *microphyllum* | 32.50±2.08 | 42.91±3.27 | -14.72** |
| *Jasminum nudiflorum* | 30.8±4.87 | 45.65±5.08 | -11.56** |
| *Jasminum officinale* | 38.37±3.86 | 48.30±5.20 | -8.40** |
| *Jasminum pentaneurum* | 36.11±2.54 | 43.15±3.75 | -8.51** |
| *Jasminum yuanjiangense* | 31.87±4.26 | 42.41±4.46 | -15.09** |

Negative t-values indicate that polar axes of S-morph are longer than those of L-morph. *: *p* < 0.05, **: *p* < 0.01.

**Table S13 Comparisons of equatorial axes of pollen between L-morph and S-morph.**

| Species | L-Morph X±SD (μm) | S-Morph X±SD (μm) | t-value |
| --- | --- | --- | --- |
| *Chrysojasminum humile* var. *microphyllum* | 32.52±0.50 | 42.10±3.35 | -14.68** |
| *Jasminum nudiflorum* | 30.75±2.42 | 43.88±4.9 | -13.16** |
| *Jasminum officinale* | 35.87±3.42 | 46.33±3.79 | -7.53** |
| *Jasminum pentaneurum* | 34.21±2.61 | 43.57±2.71 | -6.52** |
| *Jasminum yuanjiangense* | 30.47±3.88 | 40.44±3.82 | -12.60** |

Negative t-values indicate that equatorial axes of S-morph are longer than those of L-morph. *: *p* < 0.05, **: *p* < 0.01.

**Table S14 Comparisons of lumen areas between pollen from L‑morph and S‑morph.**

| Species | L-Morph X±SD (µm^2) | S-Morph X±SD (µm^2) | t-value |
| --- | --- | --- | --- |
| *Chrysojasminum humile* var. *microphyllum* | 0.87±0.38 | 1.41±0.47 | -4.93** |
| *Jasminum nudiflorum* | 2.18±0.81 | 2.1±0.43 | 1.59 |
| *Jasminum officinale* | 12.02±4.23 | 9.56±3.58 | 1.74 |
| *Jasminum pentaneurum* | 1.90±0.58 | 2.26±0.96 | -0.19 |
| *Jasminum yuanjiangense* | 2.18±0.66 | 3.64±1.33 | -5.35** |

Positive/negative t-values indicate that lumen areas of S-morph are smaller/larger than those of L-morph. *: *p* < 0.05, **: *p* < 0.01.
